# Single-cell quantitative trait loci analyses identify expression changes in macular degeneration risk genes

**DOI:** 10.64898/2026.03.30.714946

**Authors:** Andrew P. Voigt, Nathaniel K. Mullin, Kelly Mulfaul, Lola P. Lozano, Emma M. Navratil, Miles J. Flamme-Wiese, Jeremy A. Lavine, John H. Fingert, Budd A. Tucker, Edwin M. Stone, Todd E. Scheetz, Robert F. Mullins

## Abstract

Age-related macular degeneration (AMD) is a common, complex disease affecting older individuals that can lead to severe vision loss. It is characterized by early anatomical changes in the retina, retinal pigment epithelium (RPE), and choroid, especially in the central (macular) region. AMD can progress to severe atrophy and/or pathologic angiogenesis that leads to visual decline. Over 30 genetic loci have been identified as contributing to AMD risk; however, the mechanisms by which genetic variants affect pathology has not been thoroughly explored. In this report we examined single-nucleus gene expression in the retina, RPE and choroid of 88 individuals categorized by AMD stage, as well as 36 previously published samples. Genotyping was performed on 1.8 million SNPs, with additional SNPs imputed, on each donor to identify expression quantitative trait loci (eQTLs). We found that two AMD-risk loci (PILRB and ARMS2/HTRA1) affected the expression of *PILRB* and *HTRA1*, respectively. The risk allele of PILRB was associated with increased *PILRB* RNA in cones, fibroblasts, choroidal resident macrophages, and RPE, whereas the HTRA1 risk locus was associated with decreased *HTRA1* RNA in the RPE. We also identified an age-related decrease in complement inhibitors in the choriocapillaris, a tissue susceptible to complement mediated damage in AMD. We have made the eQTL and gene expression data fully available on the web resource Spectacle for rapid and interactive data access.

## INTRODUCTION

Age-related macular degeneration (AMD), a common cause of global blindness, is characterized by dysfunction of multiple tissues across the posterior pole of the human eye. Photoreceptor cells in the central retina (termed the macula) degenerate, in part due to disease across two metabolically and structurally supportive tissues called the retinal pigment epithelium (RPE) and the choroid. In AMD, vision is ultimately lost due to either atrophic changes in the photoreceptors, RPE and choroid, or from pathologic angiogenesis resulting in exudation, intraretinal fluid, and damage to the same tissues. These two forms of advanced AMD are referred to as “dry” and “wet” respectively, and the molecular drivers of progression to either stage are not well characterized.

Numerous environmental and genetic risk factors have been implicated in the development of AMD, and genome-wide association studies (GWAS) have identified 34 genetic variants associated with AMD risk^1^. The strongest genetic risk factors for AMD reside on chromosome 1 (the locus harboring the complement factor H (*CFH*) gene and its paralogs) and the 10q26 locus near genes PLEKHA1, ARMS2, and HTRA1. The identification of the CFH risk variant, which results in a Tyr402His substitution in the CFH protein, has significantly advanced our understanding of AMD pathophysiology. This variant has been shown by multiple studies to increase the deposition of monomeric C-reactive protein in the choriocapillaris and promote expression of cytokines and adhesion molecules. In contrast, the mechanism(s) by which the 10q26 risk variant results in AMD are unclear^2^. The rs10490924 variant results in an Ala69Ser substitution in ARMS2, however ARMS2 has an unknown function and has negligible RNA and protein expression in the adult retina, RPE, and choroid.

We and others believe that some genetic variants influence gene expression in the retina, retinal pigment epithelium (RPE), or choroid. Expression quantitative trait loci (eQTL) analysis is a statistical technique that can identify genetic regulators of gene expression. Identifying eQTLs in the retina and RPE has been explored by the vision research community to improve mechanistic understanding of AMD risk. For example, eQTLs have been identified for large cohorts of retinal^2^, RPE/choroid^3^, and iPSC-derived RPE cell lines derived from AMD and control donors^4^, however such studies have quantified gene expression at the bulk RNA level or only in a single cell type. Single-cell gene expression studies now permit cell-type-specific eQTL analyses, allowing for the identification of more specific regulatory mechanisms.

To improve our understanding of genetic regulation of gene expression across retinal, RPE, and choroidal cell populations, we genotyped and phenotyped postmortem human donor maculae with and without AMD in a single-cell eQTL analysis. This analysis includes genotype and single-cell gene expression from a total of 122 human donor maculae spanning controls, dry AMD, GA, and MNV. We identify cis-eQTLs across each cell type, including a decrease in *HTRA1* expression in the RPE with high-risk variants at the 10q26 locus and increased *PILRB* expression across multiple cell types with AMD risk variants near the *PILRB* gene^1^. We analyzed these data in conjunction with previous GWAS studies to identify regulatory mechanisms of AMD risk genes and gene expression changes across different aging and AMD disease states.

## RESULTS

### Single-Cell Gene Expression and Genotyping of Human Retinal, RPE, and Choroidal Cells

To identify cell-type-specific eQTLs, we generated a large single-nuclei RNA sequencing (snRNA-seq) gene expression dataset from 88 genotyped human donors spanning 89 maculae (one donor had one eye with macular neovascularization (MNV) and the other with geographic atrophy (GA), and both eyes were included). In addition, we generated genotypes from 36 of our previously sequenced scRNA-seq samples from the retina^5–7^ and RPE/choroid^8–10^. In total, 122 donors met inclusion criteria for a cell-specific eQTL analysis (see methods). The newly generated snRNA-seq dataset was created from frozen human macular donor punches of RPE and choroid (**Figure 1A**) (**SI Table 1**).

**Figure 1:**
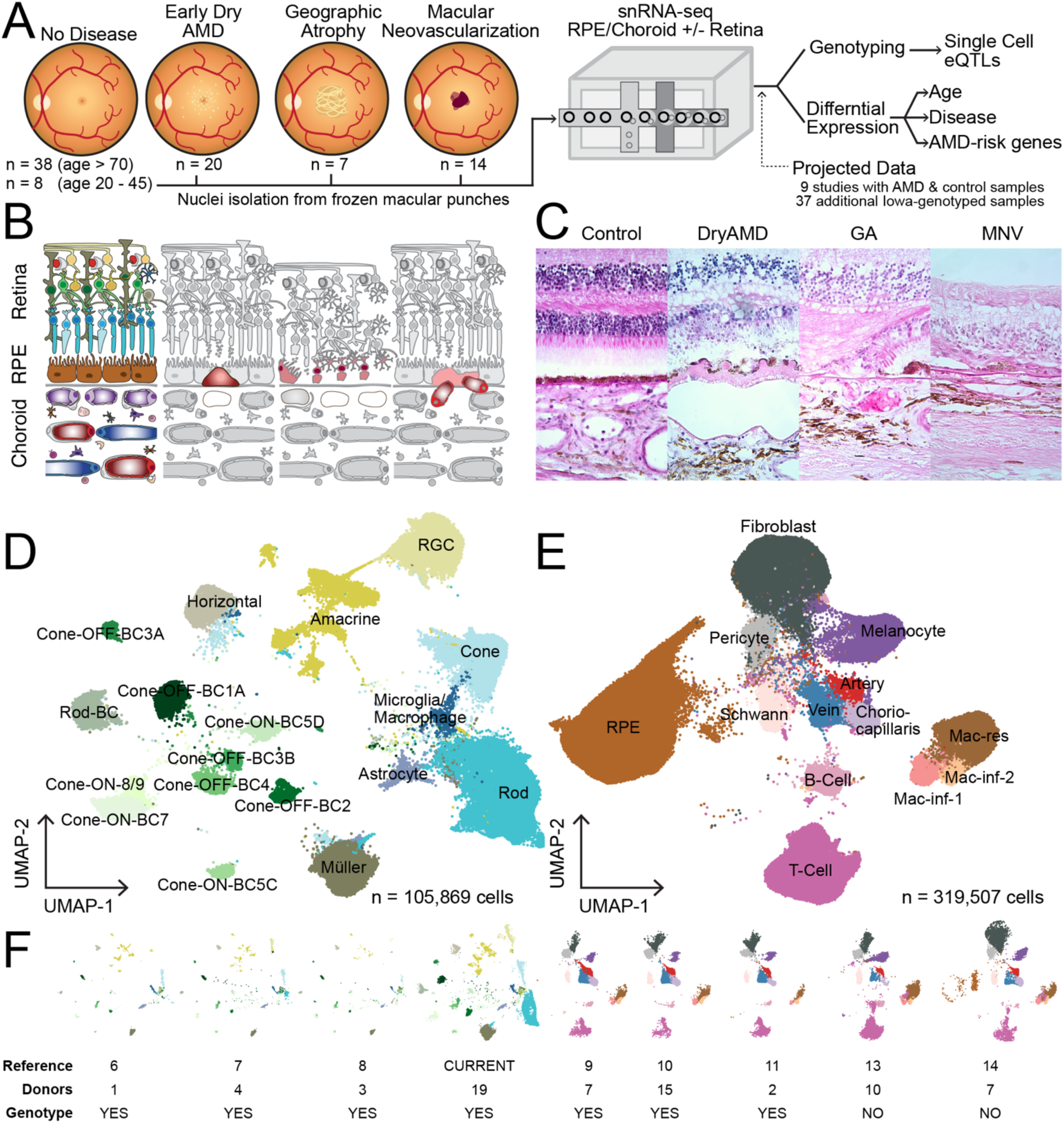
snRNA-sequencing of retina, RPE, and choroid from control and macular degeneration human donor eyes. **A.** Nuclei from 89 human donor maculae were isolated for snRNA-seq. **B-C.** Histology and medical records were reviewed and donor eyes were divided into no disease, early dry AMD, geographic atrophy, and macular neovascularization groups. Nuclei were clustered and gene expression profiles were compared to previously described retinal (**D**) and RPE/choroid (**E**) cell types. **F.** Existing human scRNA-sequencing experiments that included both control and macular degeneration donors were projected onto the dimensionality reduction of the current study for comparison of AMD-enriched genes. Previously published samples from the Iowa collection were genotyped in this study for eQTL analysis.

Neural retina tissue was recovered in 48 of these RPE/Choroidal punches, which may not be surprising as trephine biopsies can cause retinal tissue to adhere to the underlying RPE at the peripheral edges of the punch^11^. For these fortuitous samples, retinal gene expression was also analyzed. Except for one case (Eye #87), all eyes were processed within 8 hours of death with an average time to processing of 5 hours 47 minutes. Both ophthalmologic medical history and histological images (**Figure 1B-C**) were reviewed to categorize the AMD disease status of each donor. In total, we included 46 control donors, 41 AMD (20 dry AMD, 7 GA, and 14 MNV) donors, and 2 donors with other forms of retinal disease (**SI Table 1**).

The cell identity of each nucleus was classified using automated label transfer from previously published datasets^6, 9^. To better classify closely related cell subtypes, cells predicted to originate from the retina (**Figure 1D**) were separated from RPE/choroidal cell types (**Figure 1E**) before re-clustering and final classification. A total of 425,376 nuclei were recovered in this experiment (n = 105,869 from retina, n = 319,507 from RPE/choroid) and all expected major cell types were identified (**SI Figure 1**). Including the previously published scRNA-seq datasets^5–10, 12, 13^ as well as an unpublished dataset of 19 scRNA-seq libraries from human macular retina (**Figure 1F**), 105 donors were included in the RPE/choroid eQTL analysis while 74 donors were included in the retina eQTL analysis (57 shared donors had sufficient cells for at least one cell type in both tissues).

We obtained a genome-wide set of 1.8 million genotypes from each donor using the Illumina Genome Diversity Array. Additionally, we imputed 6.3 million genotypes that were in linkage disequilibrium (r^2^ greater than 0.9) with a genotyped SNP. Only donors with at least 10 cells in a respective cell type were included for eQTL analysis, and the number of donors that qualified for eQTL analysis in each cell type is provided in **SI Table 2**. Expression data were aggregated and the single-cell eQTL analysis was performed with the MatrixEQTL package adjusting for covariates including probabilistic estimation of expression residuals (PEER) factors, sex, experiment, and gene expression principal components as previously described^14^.

### Identification of single-cell eQTLs

We identified a total of 9,136 unique typed and 100,724 unique imputed eQTL single-nucleotide polymorphisms (eSNPs) across 10,786 unique genes with at least one eQTL (eGenes) that met genome wide statistical significance (FDR < 0.05) (**Table 1**). A list of all typed and imputed eQTLs is provided in **SI Table 3**. The number of non-imputed eQTLs varied between different cell populations; fibroblasts had the most eSNPs (n = 1217) while Rod Bipolar cells had the least (n = 146) (**Figure 2A**). The number of eSNPs was loosely correlated with both the number of donors included in the analysis as well as the number of total cells recovered from each cell type (**SI Figure 2**).

**Table 1:** Enumeration of eGenes (genes with at least one eQTL) and eSNPs (eQTL single-nucleotide polymorphisms) across retina and RPE/choroid single cell populations. This analysis was limited to genes that were expressed by 1% or more of cells in the cell type of interest.

|  | eGenes |  |  | eSNPs |  |  |
| --- | --- | --- | --- | --- | --- | --- |
|  | Retina | RPE/Chor | Both | Retina | RPE/Chor | Both |
| Genotyped | 1000 | 1905 | 1622 | 3307 | 5573 | 256 |
| Imputed | 1898 | 3392 | 969 | 35,687 | 61,593 | 3444 |

**Figure 2:**
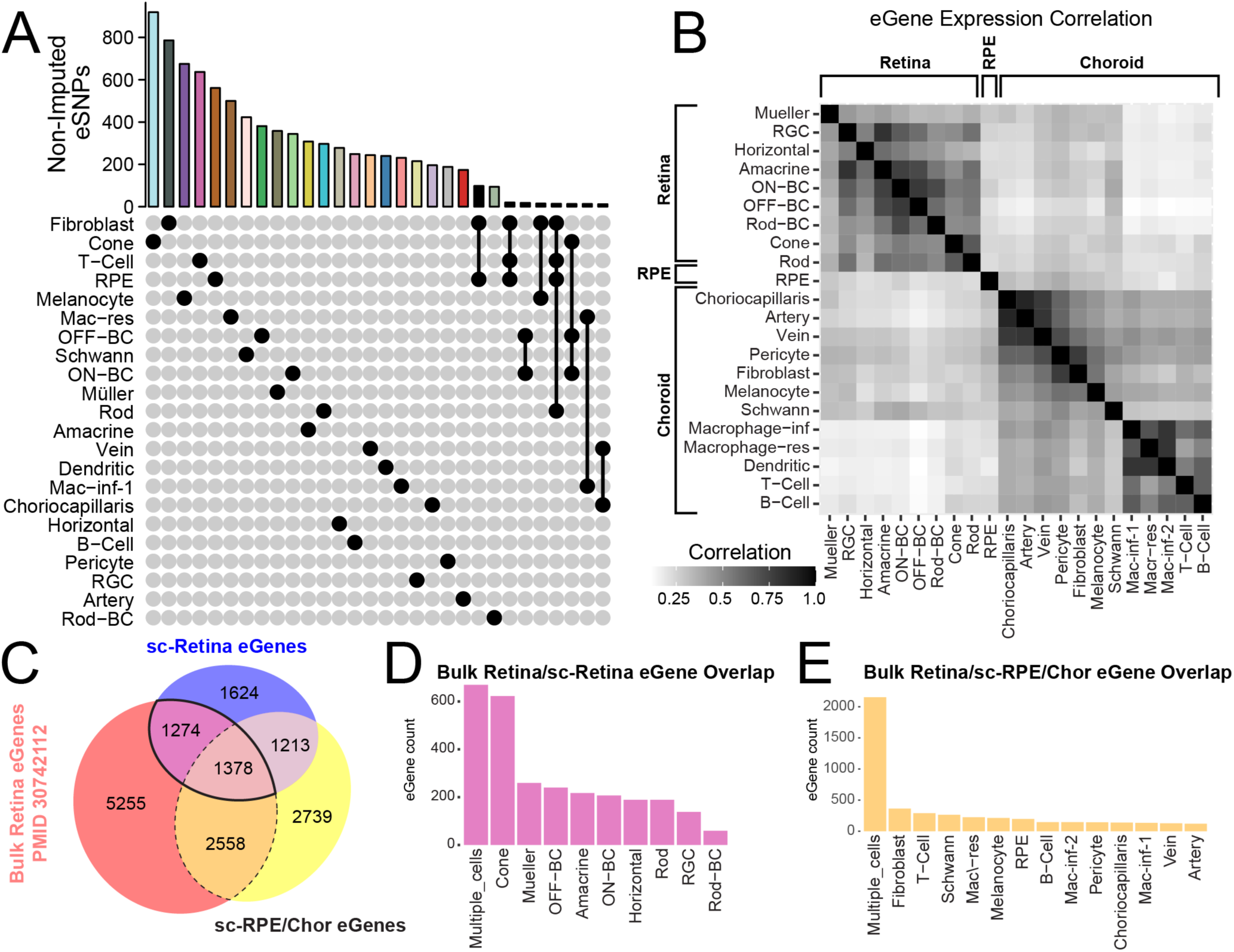
eSNP distribution across ocular cell types. **A.** Upset plot representing the number of non-imputed cell-type-specific eSNPs. Any eSNPs shared across multiple cell types are represented with vertical lines. The x-axis was truncated at 12 eSNPs in each category. **B.** For the set of non-imputed eGenes, expression was correlated across cell type 1 (x-axis) with cell type 2 (y-axis). **C.** 10,474 previously identified eGenes in a bulk retinal eQTL study^2^ (red) were compared with eGenes in the single-cell retina (blue) and RPE/choroid (yellow) cohorts. The cell types in which the overlapping eGenes met FDR significance were enumerated in the retina (**D**, containing eGenes from the solid line in panel C) and RPE/choroid (**E**, containing eGenes from the dotted line in panel C).

Of the 4527 eGenes with at least one non-imputed eSNP, we identified that 1632 (36.1%) were specific to a single cell type. As described by Yazar et al in a single-cell eQTL study of immune cells, eQTLs may be cell-specific if (1) the gene is only expressed by a single cell type, (2) there is not enough statistical power to detect an eQTL in other cell types, or (3) there are true regulatory differences across cell types^15^. To evaluate such possibilities, we first analyzed the expression of the 4507 eGenes with at least one non-imputed eSNP. We identified that only 204 eGenes (4.5%) were expressed in only one cell type. On average, 4.7 cell types expressed each eGene and 20% of eGenes were expressed by 10 or more cell types. We next asked if expression of eGenes was correlated between pairs of cell types (**Figure 2B**). Retinal cells had higher eGene expression correlation with other retinal cells compared to RPE or choroidal cells, and the same pattern was observed for choroidal cells. Indeed, more closely related cell types had more highly correlated gene expression. For example, choriocapillaris endothelial cell expression of eGenes was highly correlated with arteries (mean correlation 0.86) and veins (mean correlation 0.78), while the mean correlation for all other choroidal cell types was 0.42, with RPE cells was 0.30, and with retinal cells was 0.21.

Likewise, immune cells (macrophage, B-cell, and T-cell clusters) had more highly correlated eGene expression compared to other cell types. The correlation values observed in our study are lower than a previously published single-cell eQTL dataset of human immune cells^15^, which may be due to the more diverse cellular functions and embryonic origins of the retina, RPE, and choroidal cells. We thus conclude that the majority of eGenes are not driven by cell-specific expression but rather unique regulatory mechanisms in different cell types.

We next assessed the degree of overlap between eGenes identified by the largest previously published bulk retinal eQTL study^2^ and the current single-cell analysis. Of the 10,474 previously identified bulk retinal eQTL eGenes, 5210 met FDR significance in the current cell-specific study in at least one retinal, RPE, or choroidal cell population (**Figure 2C**). A total of 2652 overlapping eGenes were identified between the bulk study and the current single-cell retina analysis (**Figure 2C**, solid line), and 71.3% of these eGenes only met FDR significance in a single retinal cell type (**Figure 2D**). In contrast, of the 3936 overlapping bulk retina and single cell RPE/choroidal eGenes (**Figure 2C**, dashed line), 46% met FDR significance across two or more RPE/choroidal cell types (**Figure 2E**). As the neural retina and RPE/choroid have distinct embryologic origins and physiologic functions, it is perhaps not surprising that many of the previously described bulk retinal eGenes found in the current RPE/choroid population would be found across multiple RPE/choroidal cell types, perhaps reflecting shared regulatory mechanisms for this subset of eSNPs.

### eQTL analysis of AMD Risk Alleles

Previous GWAS studies identified 34 genetic variants associated with increased AMD risk^1^, and we next determined if any such variants had statistically significant single-cell eSNPs. We identified six statistically significant expression associations with these AMD risk variants (**Table 2**). Of highest interest was rs3750846, the greatest genetic risk factor for AMD, which resides on the 10q26 locus near the genes *PLEKHA1*, *ARMS2*, and *HTRA1*. The rs3750846 variant was associated with significantly decreased RPE cell *HTRA1* expression (p = 7.1×10^-7^, FDR = 0.03) (**Figure 3A-C**). This agrees with a recent large qtPCR study of a large cohort of human donors, which identified that donors with the C/C risk genotype had significantly decreased *HTRA1* mRNA in RPE cells but not bulk retinal or bulk choroid tissue^16^. Interestingly, retinal horizontal cells had the highest *HTRA1* expression of any cell type, and the rs3750846 variant had a trend towards decreased horizontal cell *HTRA1* expression without reaching statistical significance (p = 1.5×10^-4^, FDR = 0.18). We queried a previously published human single-cell ATAC sequencing experiment of retinal and choroidal tissue^17^ and observed open chromatin surrounding the rs3750846 variant in a subset of retinal cells (horizontal, amacrine, bipolar, rod photoreceptor, and cone photoreceptor cells) and in RPE cells but not in any choroidal cells (**Figure 3D**), which supports the hypothesis that this locus could be mechanistically available to participate in gene expression regulation. The surrounding genes PLEKHA1 and ARMS2 did not demonstrate expression changes with rs3750846 across any retinal, RPE, or choroidal cell type. Collectively, these results suggest that the rs3750846 variant regulates *HTRA1* expression in RPE cells and possibly horizontal cells, resulting in decreased *HTRA1* expression.

**Table 2:** eSNPs corresponding to previously identified AMD susceptibility loci.

| eSNP | AMD GWAS SNP | Gene | P-value | FDR | Cell Type |
| --- | --- | --- | --- | --- | --- |
| chr7:100393925:C:T | rs7803454 | C7orf61 | 2.79E-06 | 0.0138 | RPE |
| chr10:122456049:T:C | rs3750846 | HTRA1 | 2.44E-06 | 0.0279 | RPE |
| chr7:100393925:C:T | rs7803454 | PILRB | 9.38E-11 | 3.17E-06 | RPE |
| chr7:100393925:C:T | rs7803454 | PILRB | 4.62E-09 | 4.13E-05 | Fibroblast |
| chr7:100393925:C:T | rs7803454 | PILRB | 7.93E-09 | 1.45E-04 | Cone |
| chr7:100393925:C:T | rs7803454 | PILRB | 1.46E-06 | 0.0144 | Mac-res |

**Figure 3:**
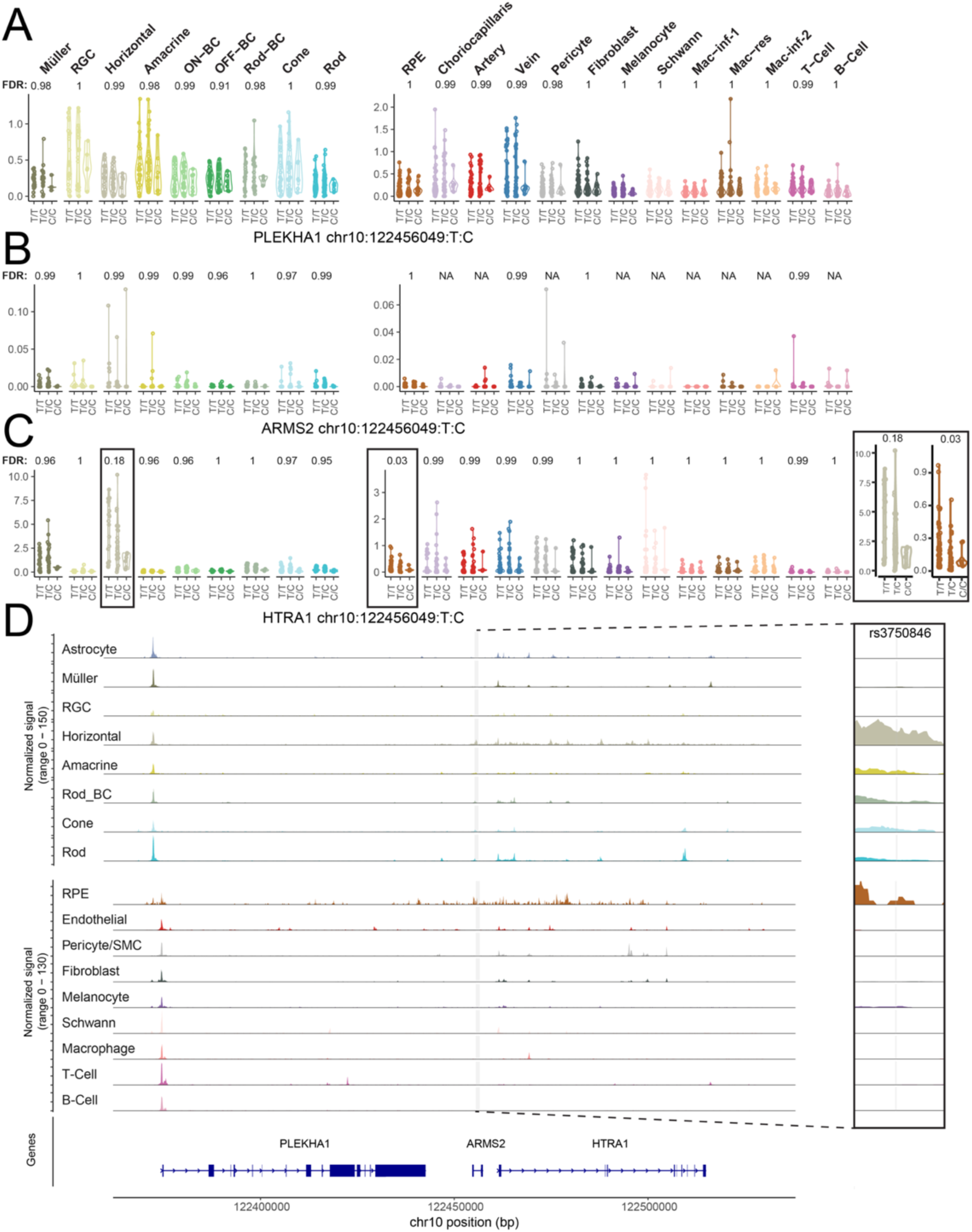
Gene expression and chromatin accessibility at the 10q26 AMD susceptibility locus. Allelic expression of *PLEKHA1* (**A**), *ARMS2* (**B**), and *HTRA1* (**C**) as a function of genotype at the rs3750846 locus. RPE cells demonstrated decreased *HTRA1* expression with increasing numbers of the C risk allele (FDR =. 0.03), and horizontal cells mirrored this trend approaching genome significance (FDR = 0.18) (insert). **D.** Chromatin accessibility at rs3750846 shows open chromatin in horizontal, amacrine, rod-bipolar, photoreceptor, and RPE cells but not choroidal cells.

rs7803454 was the only other AMD-risk variant that met genome wide statistical significance and had five identified eSNPs. The rs7803454 variant was associated with increased *PILRB* expression across several cell types (including cone photoreceptors, RPE, choroidal resident macrophages, and fibroblast cells) as well as increased expression of nearby *C7orf61* (also known as *SPACDR*) in RPE cells (**Table 2, SI Figure 3**). *PILRB* encodes the immunoglobulin-like receptor 2B, which has been implicated in the activation of the immune response^18^ and has been recently associated with photoreceptor dysfunction^19^. These genes were overall expressed at low levels across all queried cell types.

To facilitate exploration of single-cell level eQTLs, we created an interactive web browser on Spectacle^20^, available at https://singlecell-eye.org. Users can explore eQTL results and generate the genotype-expression plots produced in this manuscript for any gene or SNP of interest.

### Differential Gene Expression Across Aging and AMD States

In addition to identifying genetic variants implicated in gene expression regulation, we next identified genes enriched in different AMD states by performing a pseudobulk differential expression analysis between control cells and each of early dry AMD, geographic atrophy, and MNV donors (**SI Files 4-7**), with filtered results for more highly expressed genes in each cell type (**SI Files 8-11**, see methods). To better understand how known AMD risk genes contribute to disease pathogenesis, we first analyzed differential expression results of known AMD risk genes across dry AMD, MNV, and GA samples (**Figure 4**). AMD risk genes were either identified in previous GWAS studies^1^ or are members of the GSEA complement gene set, as complement is a major risk factor of AMD^21^. To focus on gene expression changes that are most likely to be biologically meaningful, we first quantified the relative expression of each AMD-associated gene across all cell types (**Figure 4**, white-to-black color scale), and next focused the differential expression analysis (blue-to-red color scale) on cells that express a gene at 33% or more of the maximum value.

**Figure 4:**
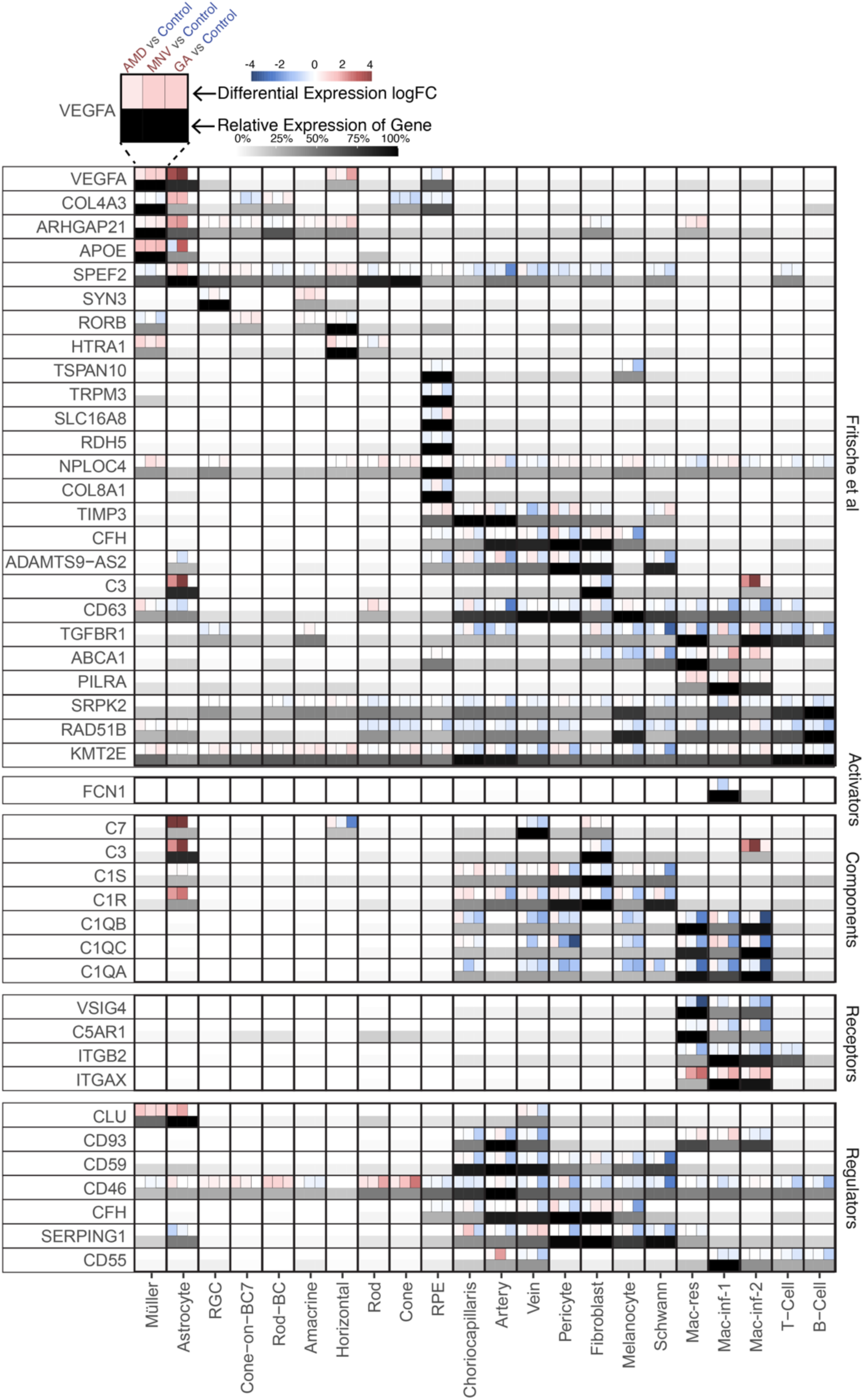
Differential expression of AMD risk genes across disease states. For each AMD risk gene, the relative expression in each cell type is shown in a white-to-black color gradient. Above the relative expression, the log2FC for each AMD comparison is shown with dry AMD on the left, MNV in the center, and GA right. Blue colors indicate enrichment in control samples and red colors indicate enrichment in AMD states.

The retinal macroglial cells (Müller glia and astrocytes) highly and differentially expressed several AMD risk genes. Müller glia and astrocyte *VEGFA* expression, which encodes an angiogenic growth factor that also promotes vascular permeability^22^, was increased across dry AMD (left most block), MNV (middle block), and GA (right most block) samples (**Figure 4**). Likewise, expression of apolipoprotein E (*APOE*), a major regulator of cholesterol and lipid handling in the retina, was upregulated across all three AMD states in Müller glial cells. Several complement genes were differentially expressed across AMD states. For example, AMD-associated astrocytes expressed higher levels of complement components *C1S*, *C1R*, *C3*, and *C7* compared to control samples. Similarly, choroidal macrophages in AMD samples had increased expression of *ITGAX*, a gene that encodes CD11c and is a marker of proangiogenic macrophages^23^. Lastly, many complement inhibitors, including *CD46*, *CD59*, and *CFH,* demonstrated decreased expression in AMD-associated choroidal endothelial and stromal cells, supporting the hypothesis that complement is over-activated in AMD choroids.

As age is the greatest risk factor for developing AMD, we next explored how expression of AMD risk genes changes with donor age. Our snRNA-seq cohort included 8 “young” control donors less than 42 years old (mean 32.2 years old, range 22 - 42 years old) as well as 38 control donors greater than 70 years old (mean 84.1 years old, range 70 - 97 years old). We completed a pseudobulk differential expression analysis between these young and aged control cohorts for all retinal, choroidal, and RPE cell populations (**SI File 5**). First, we restricted our analysis to known AMD risk genes (**Figure 5**).

**Figure 5:**
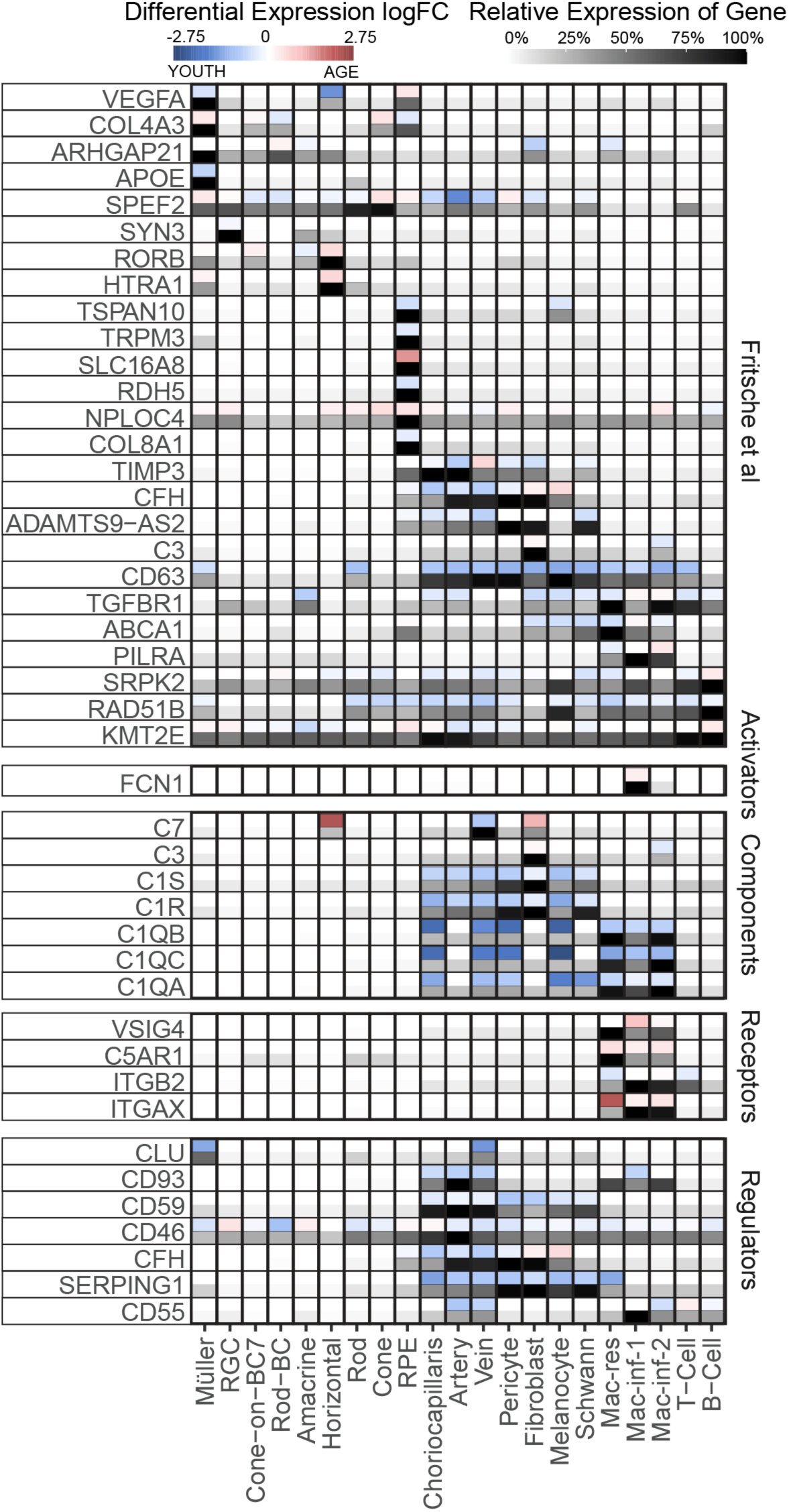
Differential expression of AMD risk genes between young and aged cohorts. For each AMD risk gene, the relative expression in each cell type is shown in a white-to-black color gradient. Above the relative expression, the log2FC for the young (blue) versus aged (red) cohorts is depicted.

Interestingly, most AMD-risk genes demonstrated decreased expression in the aging patient cohort. Both complement components (such as *C1QA*, *C1QB*, and *C1QC*) and regulators (such as *CFH*, *CD46*, and *CD59*) showed decreasing expression with advancing age, supporting the hypothesis that complement is dysregulated in aging. While the aged cohort had relatively fewer gene enrichments, aged RPE cells demonstrated higher expression of *SLC16A8*, a monocarboxylate transporter 3 which transports lactate from the outer retina to the choroid^24^, and aged macrophages were slightly enriched in the complement receptor *C5AR1*.

Lastly, we identified novel differentially expressed genes without previous direct genetic association to AMD. Given the large number of comparisons, we focused our discussion to three cell types central to AMD pathogenesis: rod photoreceptor cells, RPE cells, and choriocapillaris endothelial cells, however differential expression results for all cell types are available (**SI Files 2-5**). For these cell types, we display the 10 most differentially expressed genes in each direction for the AMD versus control, MNV versus control, and youth versus aged control cohorts from the filtered differential expression lists (**Figure 6**). In addition, we compare the fold changes quantified in this study with previously published scRNA-seq experiments (see **Figure 1F**).

**Figure 6:**
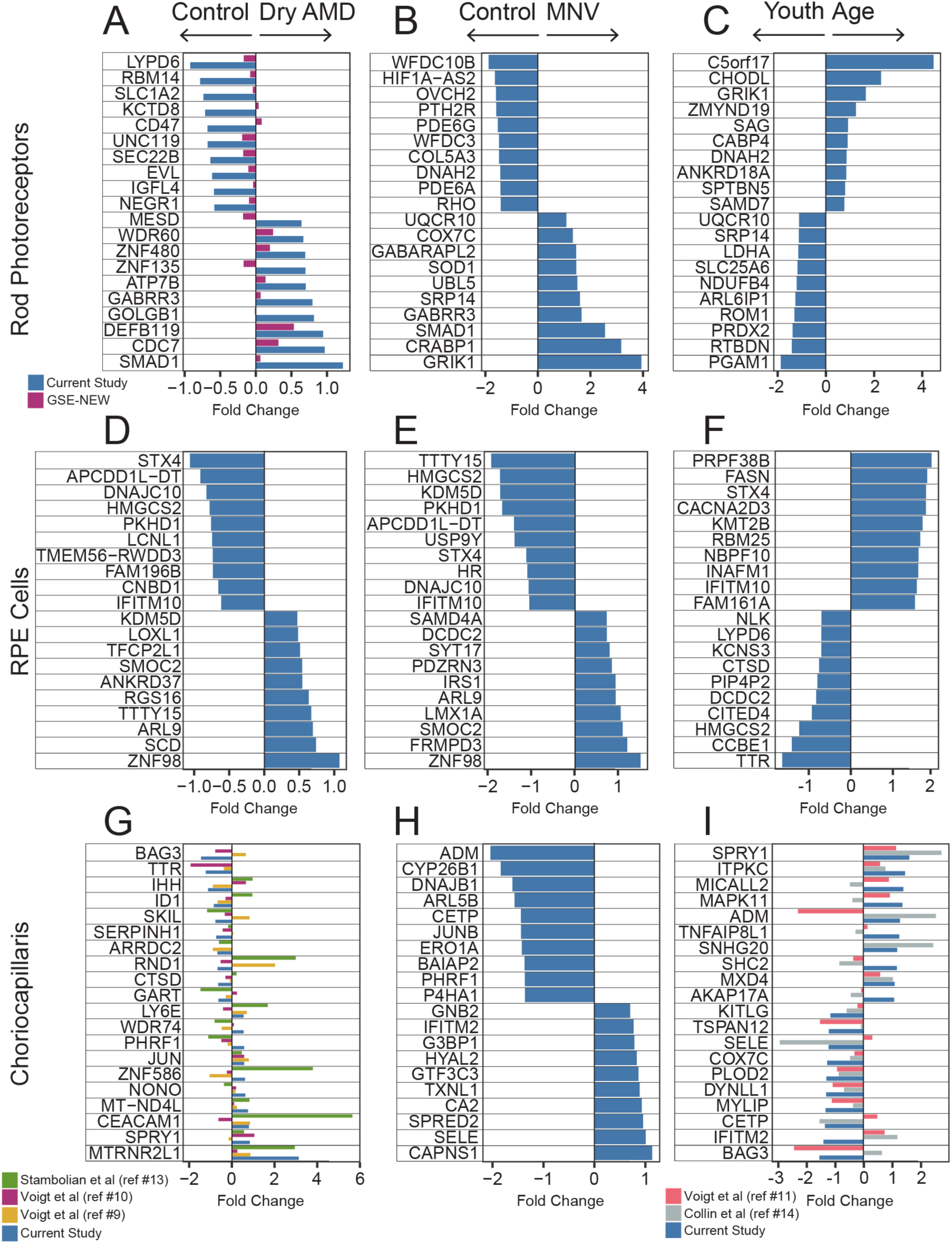
Top Differential Expression Results for Rod Photoreceptors, RPE Cells, and Choriocapillaris endothelial cells across Dry AMD, MNV, and Age. The top and bottom 10 differentially expressed genes for rod photoreceptors (**A-C**), RPE cells (**D-F**), and choriocapillaris endothelial cells (**G-I**) are displayed for dry AMD versus control (column 1), MNV versus control (column 2), and young versus aged cohorts (column 3). When possible, differential expression results from previously published single-cell RNA sequencing studies were compared to log fold changes in the current study.

Rod photoreceptors are the first neural retinal cell type to degenerate in early AMD^25^, and we first examined the most differentially expressed genes in rods across AMD, MNV, and age (**Figure 6A-C**). The most AMD-enriched rod photoreceptor gene was *SMAD1*, which encodes a bone morphogenic protein signal transducer which has been shown to increase after retinal damage^26^ (**Figure 6A**).

Other AMD and control enriched rod gene expression changes were highly concordant with the newly generated single-cell RNA sequencing study of 11 Control versus 8 AMD donors (**Figure 6A**). In the aged analysis, rod photoreceptors from aging donors lost expression of *RTBDN*, a riboflavin-binding protein which has been implicated in photoreceptor protection during retinal degeneration^27^. Likewise, rod photoreceptors from the aged cohort had decreased *ROM1*, which is necessary for rod photoreceptor viability^28^ and *ARL6IP1,* an anti-apoptosis protein^29^. These results suggest that rod photoreceptors from aged donors may decrease expression of genes implicated in structural stability and cell viability.

Located directly beneath photoreceptor cells, RPE cells play a crucial role in retinal homeostasis and dysfunction in early AMD. Both control and young donors demonstrated higher expression of *HMGCS2 (***Figure 6D-F**), a ketogenic enzyme that is involved in utilizing photoreceptor outer segments for metabolism^30^. RPE cells from MNV samples had increased expression of *SMOC2*, a proangiogenic factor that potentiates VEGF signaling^31^. In the aging analysis, the aged cohort lost expression of transthyretin (*TTR)*, a vitamin A transporter^32^, while older RPE cells expressed increased fatty acid synthase (*FASN)*, which is associated with abnormal lipid accumulation^33^.

Choriocapillaris endothelial cell loss is among the first histopathological hallmarks of AMD^34^. *MTRNR2L1* was the most upregulated gene in the choriocapillaris AMD analysis (**Figure 6G**) and was also increased in AMD samples across three previous single-cell studies. *MTRNR2L1* encodes a pseudogene for the mitochondrially-derived peptide humanin, and interestingly, *MTRNR2L1* was upregulated in AMD across other retinal and choroidal cell types (**SI Figure 4**). While enrichment across multiple distinct cell types is suspicious for contamination or dying cells, *MTRNR2L1* was not upregulated in the geographic atrophy or MNV donors (**SI Figure 4**) nor did cells from AMD donors have increased damage-associated gene expression hallmarks of mitochondrial or ribosomal gene contamination, suggesting that *MTRNR2L1* expression may be associated with AMD^35^. Both aging and dry AMD choriocapillaris endothelial cells had increased expression of *SPRY1*, a mediator of cellular senescence that is upregulated in hypoxia^36^ (**Figure 6G-I**). In contrast, both control and young choriocapillaris endothelial cells had increased expression of *BAG3,* which promotes endothelial cell survival and stress responses^37^. MNV-derived choriocapillaris endothelial cells demonstrated increased expression of calpain small subunit 1 (CAPNS1), which stabilizes the proangiogenic molecules calpain 1 and 2^38^ and the proangiogenic adhesion molecule e-selectin (SELE)^39^ (**Figure 6H**).

## DISCUSSION

In heterogenous diseases such as AMD, genotype-phenotype correlations provide insight into genetic regulation of disease-associated traits. In this study, we expand on prior bulk eQTL analyses of ocular tissues^2–4^ and perform a large cell-type-specific eQTL analysis across human retinal, RPE, and choroidal cell populations. This analysis was completed across 122 post-mortem donor eyes and identifies cell-type-specific eQTLs not detected by prior bulk studies. Additionally, we show that rs3750846, the polymorphism associated with high AMD risk^1^, is associated with a significant decrease in *HTRA1* gene expression in RPE cells. Finally, differential gene expression across dry AMD, GA, MNV, young, and aging conditions identifies dysregulation of many genes, including components of the complement system, with age and disease.

Prior GWAS studies have identified multiple genetic risk factors for AMD, including non-coding variants in the 10q26 locus^1^. This locus contains three genes, *PLEKHA1*, *ARMS2*, and *HTRA1,* that reside in proximity, and no consensus has been reached regarding which gene(s) or protein(s) at this locus are affected by the rs3750846 polymorphism and mechanistically lead to AMD. Previous studies using retina^40^ or RPE/choroid^41^ showed no difference in *HTRA1* expression with genotype, although sample sizes were small (n = 35 and 82, respectively) and used bulk RNA, which may limit the sensitivity to detect cell-specific gene expression changes. In circulating monocytes derived from peripheral blood, the high-risk 10q26 haplotype demonstrated increased *HTRA1*, and it has been purported that such HTRA1 upregulation drives TSP1-mediated inflammation and mononuclear phagocyte persistence in the subretinal space^42^. Our single cell eQTL analysis corroborates a recent finding^16^ that RPE cells have decreased expression of *HTRA1* with increasing copies of risk alleles at rs3750846, and adds to this previous investigation by showing that this gene expression change is not present in any other retinal or choroidal cell population (**Figure 3**). However, in the retina, horizontal cells showed a trend towards rs3750846-dependent decrease in *HTRA1* expression (p = 1.5×10^-4^, FDR = 0.18), which is notable as retinal horizontal cells, bipolar cells, and photoreceptors have open chromatin surrounding rs3750846 (**Figure 3D**). Similar to Williams et al, we also observed a slight decrease in RPE *HTRA1* expression with age (log2FC = 0.42, SI Table 7), and it is possible that a corresponding decrease in HTRA1 protein could lead to impaired Bruch’s membrane maintenance and downstream AMD changes^16^.

Prior GWAS studies have also identified that variants in the paired immunoglobulin-like type 2 receptor B (PILRB) are associated with AMD, and our single-cell eQTL analysis identifies increased expression of *PILRB* with increasing risk alleles in RPE cells, fibroblasts, cone photoreceptors, and resident macrophages. PILRB has been primarily studied in the context of immune cells where it increases inflammation and activates adaptive immune cells^38^. However in gastric cancer, PILRB has been shown to increase the synthesis of cholesterol^38^, a major component of drusen. PILRB also is implicated in photoreceptor stability, and loss of PILRB in mice leads to photoreceptor ribbon synapse defects and decreased electroretinogram responses^19^. While further mechanistic studies are required to unravel the pathogenesis of such *PILRB* genomic variants, it is notable *PILRB* mRNA is increased across several cell types in the choroid, RPE, and retina with an increasing number of risk alleles.

In addition to cell-type-specific eQTLs, the snRNA-seq experiment provides gene expression data across a very large population of human donor retinas, RPE, and choroids, spanning dry AMD (n = 20), GA (n = 7), and MNV (n = 14) phenotypes. Complement overactivation is a hallmark of AMD, and each of the AMD states demonstrated dysregulated complement gene expression (**Figure 4**). We also observed an age-related decrease in expression of complement regulators such as *CFH*, *CD46*, and *CD59* particularly in choroidal endothelial and stromal cells (**Figure 5**). Such differential expression results, which are provided in their entirety for all cell types (**SI Files 4-11**), can help us better understand pathogenesis mechanisms for different AMD subtypes.

There is a growing appreciation that genetic regulation in complex disease is context and cell-type-dependent. By integrating genotypes, RNA expression, and phenotypic AMD status, we have an improved understanding of the cell-type-specific disease mechanisms of genetic drivers of AMD. To facilitate reanalysis, all raw and processed gene expression data as well as all genotype data can be downloaded at publicly available data repositories. In addition, we have built an interactive single-cell eQTL browser on the freely available Spectacle website (https://singlecell-eye.org) to assist in rapid exploration of these data.

## METHODS

### Human donor maculae

Human eyes were obtained following complete informed consent from the donors’ next of kin through the Iowa Lions Eye Bank (Iowa City, IA) and in compliance with the Declaration of Helsinki. In all cases consent included access to medical information. Only samples from deceased individuals were used in this study. Anterior segments containing the cornea, iris, and ciliary body were removed and posterior poles were dissected by making 4 equidistant, full thickness incisions from anterior to posterior, leaving the optic nerve head and macula intact. Photographs that included the macula were collected using a Leica 9si microscope to aid subsequent diagnosis. Macular and extramacular punches of neural retina and RPE/choroid were obtained and flash frozen in liquid nitrogen. For macular tissue, either a complete 6mm punch or, in most cases, a bisected 8mm punch was obtained. For these maculae the other half was fixed in 4% paraformaldehyde for histological analyses and for assessing pathology. All samples in this study were preserved within 8 hours of death. Tissues were maintained at -70 to -80 degrees until used for nuclear isolation.

Eyes were categorized as controls, early-intermediate AMD, geographic atrophy, or macular neovascularization using chart review, examination of gross photos, and/or examination of histology.

### Single-Cell RNA Sequencing

Nuclei were isolated using the 10x Genomics Chromium Nuclei Isolation kit according to the manufacturer’s instructions. Briefly, punches were removed from the freezer and immediately placed on dry ice. Samples were transferred to pre-chilled sample dissociation tubes and samples dissociated, followed by addition of lysis buffer to the sample tube. Tissue was then ground with a disposable pestle to liberate nuclei, followed by additional lysis buffer and mixing by pipetting. Tissue homogenates were filtered through a 100µm strainer to remove debris and eluant was added to prechilled nuclei isolation collection tubes. Nuclei were isolated by centrifugation at 16,000xg followed by washing and resuspending the nuclear pellet. Nuclei were separated and barcoded with the Chromium Controller instrument (10X Genomics) and Single Cell 3′ Reagent (v3.1 chemistry) kit (10X Genomics) according to the manufacturer’s specifications and as described previously^43^ in batches of 8-16 samples. snRNA libraries were pooled and sequencing was performed by the Genomics Division of the Iowa Institute of Human Genetics using the NovaSeq 6000 instrument (Illumina) generating 100 bp paired end reads. FASTQ files were generated from base calls with bcl2fastq software (Illumina).

Reads were mapped to the human genome (GRCh38) with CellRanger 7.0 Cells were filtered from downstream analysis if they had less than 550 counts, less than 450 or greater than 7000 unique RNA features or had more than 12.5% of reads mapping to mitochondrial reads. To minimize ambient RNA contamination, the R package SoupX^44^ was used to estimate and remove ambient RNA expression. After SoupX, cells were clustered and canonical marker genes were used to divide into retinal neural tissue versus RPE/choroidal tissue. Integration was performed independently on retina and RPE/choroidal samples within Seurat v5^45^ using the Harmony Integration method. The R package DoubletFinder^46^ was utilized to predict cell-cell doublets which were removed from downstream analysis.

### Genotyping

Genomic DNA was isolated from peripheral retina of each donor using the Qiagen DNeasy Blood and Tissue kit according to the manufacturer’s instructions. A total of 200 ng of high purity DNA from each sample was ran on the Illumina Infinium Genome Diversity Array-8 v1.0 platform at the Genomics Division of the Iowa Institute of Human Genetics. Genotypes were called from the IDAT files using the Illumina Array Analysis Platform Genotyping Command Line Interface (v1.1), converted to the BCF format, filtered to remove CNV calls, and split into per-chromosome VCF files. Imputation of additional variations and phasing was performed using the Michigan Imputation Server^47^ using MiniMac4 for imputation, Eagle for phasing, an Rsq filter of 0.3, and the 1000G Phase3 Low panel. The imputed files were then concatenated into a merged BCF file and filtered to remove variants with an observed minor allele frequency of less than 5%.

### Cell-Type-Specific eQTL Analysis

Cell-type-specific eQTL analysis was completed by taking the pseudobulk expression data of each cell type and calculating PEER factors (latent variables that explain gene expression variability) as previously described^14^. Briefly, pseudobulk gene expression was calculated for each donor that had 10 or more cells recovered in the cell type of interest. A total of 10 PEER factors were calculated using the pseudobulk gene expression data with the peer R package^48^ applying 2000 maximum iterations and the first 5 gene expression principal components. The MatrixEQTL R Package^49^ was used to test for association between genotype and gene expression with linear regression after adjusting for covariates including PEER factors, sex, experiment, and gene expression principal components. Analysis was limited to SNPs in the *cis*-region within 1 Mb of the gene and with a minor allele frequency of at least 5%. Statistically significant eSNPs had an FDR < 0.05.

### Differential Expression

A pseudobulk differential expression analysis was completed in which the counts for all reads in each independent library were summed before differential expression^50^. Compared to analyses in which each cell is consider an independent biological observation, the pseudobulk comparison more accurately captures the number of biological replicates in each condition. In the differential expression analysis of known AMD risk genes and complement genes, cell types that expressed a gene at less than 33% of the maximum cell type expression were excluded from analysis. Differential expression results from all cell types are fully available in **SI Files 4-7.** Additional filtering was completed to narrow differential expression results to genes meeting three minimum expression criteria in each respective cell type: (1) the gene has an absolute expression level > 0.75 in the cell type of interest, (2) the expression of the gene in the cell type of interest is greater than 33% of the expression in the cell type with the highest expression of the gene, and (3) the gene is expressed in at least 5% of cells in the cell type of interest (**SI Files 8-11**).

## Acknowledgements

We wish to gratefully acknowledge the donors and their families who gifted ocular tissue that made these experiments possible.

## Supplementary Information

**SI Table 1:** Donors for the newly generated snRNA-seq analysis: Note Eye #72 and Eye #73 are from the same donor; with the former demonstrating geographic atrophy and the later with macular neovascularization.

| Eye | Age | Gender | Disease State | Preservation Time | Cause of Death |
| --- | --- | --- | --- | --- | --- |
| Eye 1 | 21 | Female | Control | 5:25 | Hodgkins Lymphoma |
| Eye 2 | 42 | Male | Control | 6:30 | Sepsis |
| Eye 3 | 32 | Male | Control | 6:17 | Leukemia/Cardiac Arrest |
| Eye 4 | 42 | Female | Control | 6:32 | Acute Liver and Renal Failure |
| Eye 5 | 36 | Female | Control | 5:54 | Drug Overdose |
| Eye 6 | 23 | Male | Control | 5:52 | Respiratory Failure/Testicular Cancer |
| Eye 7 | 22 | Female | Control | 6:55 | Cardiopulmonary Arrest/Pneumonia |
| Eye 8 | 40 | Male | Control | 5:40 | Unknown |
| Eye 9 | 77 | Male | Control | 5:31 | Renal Failure |
| Eye 10 | 81 | Female | Control | unknown | Unknown |
| Eye 11 | 87 | Female | Control | 5:20 | Stroke |
| Eye 12 | 89 | Female | Control | 5:14 | Acute Subdural Hematoma |
| Eye 13 | 87 | Female | Control | 5:30 | Stroke |
| Eye 14 | 90 | Male | Control | 5:49 | Cancer |
| Eye 15 | 86 | Female | Control | 5:12 | Respiratory Failure |
| Eye 16 | 88 | Male | Control | 3:05 | Unknown |
| Eye 17 | 89 | Female | Control | 3:33 | Pneumonia |
| Eye 18 | 96 | Male | Control | 5:47 | Ischemic Stroke |
| Eye 19 | 72 | Female | Control | 6:33 | Sepsis |
| Eye 20 | 70 | Male | Control | 6:02 | Unknown |
| Eye 21 | 91 | Female | Control | 7:45 | Malnutrition |
| Eye 22 | 84 | Female | Control | 5:35 | Heart Attack |
| Eye 23 | 86 | Male | Control | 4:55 | Aortic Stenosis |
| Eye 24 | 87 | Female | Control | 6:56 | Septic Shock |
| Eye 25 | 83 | Female | Control | 6:42 | Gastric Bowel Perforation |
| Eye 26 | 91 | Female | Control | 6:20 | Pulmonary Embolism |
| Eye 27 | 90 | Female | Control | 6:10 | Cholecystitis |
| Eye 28 | 71 | Female | Control | 6:30 | Liver Cirrhosis/Sepsis |
| Eye 29 | 82 | Female | Control | 5:36 | Dry Gangrene/Supratherapeutic INR & Upper GI Bleed |
| Eye 30 | 94 | Male | Control | 6:52 | Vagal Episode |
| Eye 31 | 77 | Female | Control | 6:43 | Cardiogenic Shock/Sepsis |
| Eye 32 | 97 | Female | Control | 7:08 | Sepsis/Right Femur Fracture |
| Eye 33 | 79 | Male | Control | 6:20 | Sepsis |
| Eye 34 | 85 | Male | Control | 6:43 | Cardiogenic Shock |
| Eye 35 | 92 | Male | Control | 5:45 | Natural Causes |
| Eye 36 | 84 | Male | Control | 4:07 | Unknown |
| Eye 37 | 83 | Male | Control | 5:45 | Cardiopulmonary Arrest |
| Eye 38 | 96 | Female | Control | 5:26 | COPD |
| Eye 39 | 76 | Female | Control | 5:05 | Unknown |
| Eye 40 | 92 | Female | Control | 5:02 | Respiratory Failure Secondary to Ischemic Stroke |
| Eye 41 | 89 | Male | Control | 6:10 | End Stage Renal Disease |
| Eye 42 | 70 | Male | Control | 5:52 | Intrahepatic Bile Duct Carcinoma with Mets |
| Eye 43 | 70 | Male | Control | 5:31 | Acute Hypoxic Respiratory Failure |
| Eye 44 | 78 | Male | Control | 7:37 | Heart Failure & Pneumonia |
| Eye 45 | 76 | Male | Control | 6:04 | Unknown |
| Eye 46 | 79 | Male | Control | 5:11 | Unknown |
| Eye 47 | 77 | Male | AMD | 6:04 | LLL Pneumonia |
| Eye 48 | 89 | Female | AMD | 4:24 | Cerebrovascular Accident |
| Eye 49 | 83 | Male | AMD | 5:25 | Motor Vehicle Collision |
| Eye 50 | 79 | Male | AMD | 6:47 | Pancreatic Cancer |
| Eye 51 | 92 | Female | AMD | 5:25 | Lung Cancer |
| Eye 52 | 89 | Female | AMD | 3:54 | Unknown |
| Eye 53 | 85 | Male | AMD | 6:06 | Acute Coronary Syndrome |
| Eye 54 | 89 | Male | AMD | 6:20 | CHF |
| Eye 55 | 76 | Female | AMD | 8:02 | Unknown |
| Eye 56 | 79 | Female | AMD | 6:43 | Congestive Heart Failure |
| Eye 57 | 85 | Female | AMD | 4:21 | Cardiogenic Shock |
| Eye 58 | 93 | Female | AMD | 4:31 | Cardiac Arrest |
| Eye 59 | 83 | Male | AMD | 6:52 | Motor Vehicle Accident |
| Eye 60 | 71 | Male | AMD | 4:43 | Cardiac Arrest |
| Eye 61 | 73 | Male | AMD | 6:33 | Unknown |
| Eye 62 | 86 | Male | AMD | 7:27 | COPD |
| Eye 63 | 92 | Male | AMD | 4:33 | Multi-Organ Failure, Respiratory Failure |
| Eye 64 | 96 | Female | AMD | 5:22 | Sepsis |
| Eye 65 | 89 | Female | AMD | 7:18 | Bladder Cancer with Bone Mets |
| Eye 66 | 84 | Male | AMD | 5:22 | Unknown |
| Eye 67 | 87 | Female | GA | 4:53 | Leukemia, acute renal failure |
| Eye 68 | 87 | Female | GA | 4:31 | Unknown |
| Eye 69 | 76 | Female | GA | 4:04 | Unknown/No Records |
| Eye 70 | 87 | Female | GA | 4:56 | Cardiac Arrest |
| Eye 71 | 92 | Female | GA | 5:21 | Respiratory Distress |
| Eye 72 | 93 | Female | GA | 7:31 | Cardiogenic Shock |
| Eye 73 | 93 | Female | MNV | 7:31 | Cardiogenic Shock |
| Eye 74 | 97 | Male | MNV | 3:59 | Cardiac Arrest |
| Eye 75 | 83 | Female | MNV | 4:42 | Subdural Hematoma |
| Eye 76 | 92 | Male | MNV | 4:47 | Sepsis - Pneumonia |
| Eye 77 | 77 | Male | MNV | 5:58 | Unknown |
| Eye 78 | 80 | Male | MNV | 4:35 | Respiratory Arrest |
| Eye 79 | 70 | Female | MNV | 7:50 | Cardiac Arrest |
| Eye 80 | 82 | Female | MNV | 6:11 | Acute Renal Failure |
| Eye 81 | 92 | Female | MNV | 6:43 | Sepsis |
| Eye 82 | 83 | Female | MNV | 4:51 | Metastatic Melanoma |
| Eye 83 | 87 | Female | MNV | 4:13 | Small bowel obstruction |
| Eye 84 | 81 | Male | MNV | 7:23 | Cardiac Arrest |
| Eye 85 | 72 | Female | MNV | 6:01 | Post Surgical Complications - Bowel Resection |
| Eye 86 | 85 | Female | MNV | 4:55 | Aortic Aneurysm |
| Eye 87 | 56 | Female | Contralateral eye with chorioretinal lesions | 9:28 | Cardiac Arrest |
| Eye 88 | 97 | Female | GA | 6:06 | Cerebral Vascular Accident |
| Eye 89 | 57 | Male | ABCA4 heterozygote | 4:55 | Cardiopulmonary Arrest Secondary to Ischemic Stroke and NSTEMI |

**SI Table 2:** Enumeration of total donors included in the eQTL analysis for each cell type. Donors were required to have at least 10 recovered cells within a given cell type to be included in the eQTL analysis. OFF-bipolar-cell and ON-bipolar-cell subtypes were combined to increase the number of donors available for eQTL analysis. Cell types with less than 40 donors were excluded from cell-type-specific eQTL analysis.

| # Donors | Cell Type | Tissue |
| --- | --- | --- |
| 60 | Amacrine | Retina |
| 60 | Cone | Retina |
| 49 | Horizontal | Retina |
| 50 | Mueller | Retina |
| 55 | OFF-BC | Retina |
| 56 | ON-BC | Retina |
| 49 | RGC | Retina |
| 73 | Rod | Retina |
| 48 | Rod-BC | Retina |
| 75 | Artery | Choroid |
| 59 | B-Cell | Choroid |
| 79 | Choriocapillaris | Choroid |
| 76 | Macrophage-inf-2 | Choroid |
| 103 | Fibroblast | Choroid |
| 83 | Macrophage-inf-1 | Choroid |
| 93 | Macrophage-res | Choroid |
| 92 | Melanocyte | Choroid |
| 79 | Pericyte | Choroid |
| 80 | RPE | RPE |
| 76 | Schwann | Choroid |
| 98 | T-Cell | Choroid |
| 81 | Vein | Choroid |

**SI Table 3:** All eSNPs with an FDR < 0.05, at least 1% expression in a cell of interest, and with 3 or more total minor alleles observed in the donor cohort are reported.

**SI Files 4 – 11:** Differential expression results are reported in their entirety for control versus early dry AMD (**SI File 4**), GA (**SI File 5**), MNV (**SI File 6**). The youth versus aging differential expression is reported in **SI File 7**. Additional filtering was completed to narrow differential expression results to genes meeting three minimum expression criteria in each respective cell type: (1) the gene has an absolute expression level > 0.75 in the cell type of interest, (2) the expression of the gene in the cell type of interest is greater than 33% of the expression in the cell type with the highest expression of the gene, and (3) the gene is expressed in at least 5% of cells in the cell type of interest. Such filtered files depict differential expression in control versus early dry AMD (**SI File 8**), GA (**SI File 9**), MNV (**SI File 10**), and youth versus aging (**SI File 11**).

## Supplemental Figures

**SI Figure 1:**
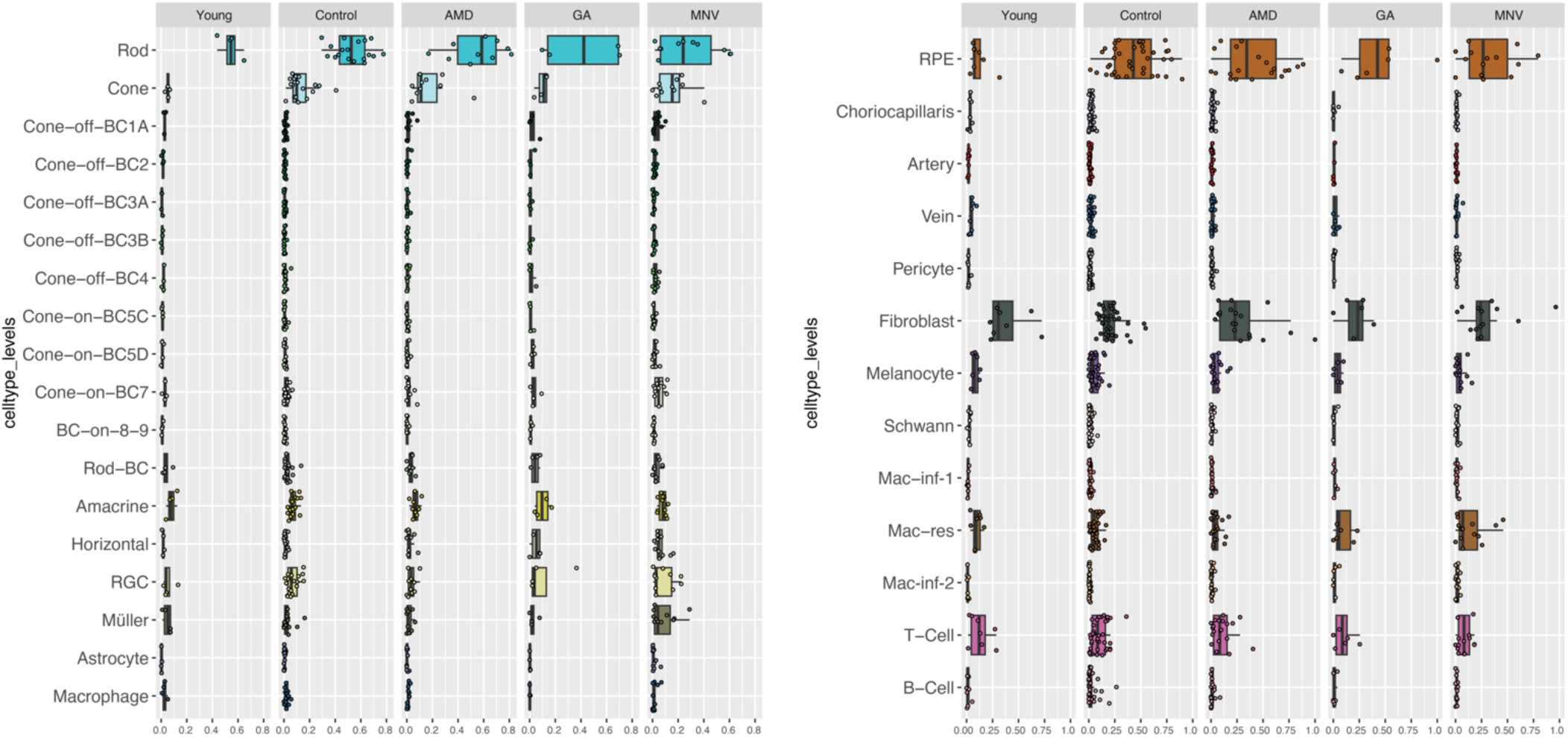
Recovered cell proportions split by disease state. Each dot corresponds to one donor, and the box plots represent the median percentage of cells classified to each cell type.

**SI Figure 2:**
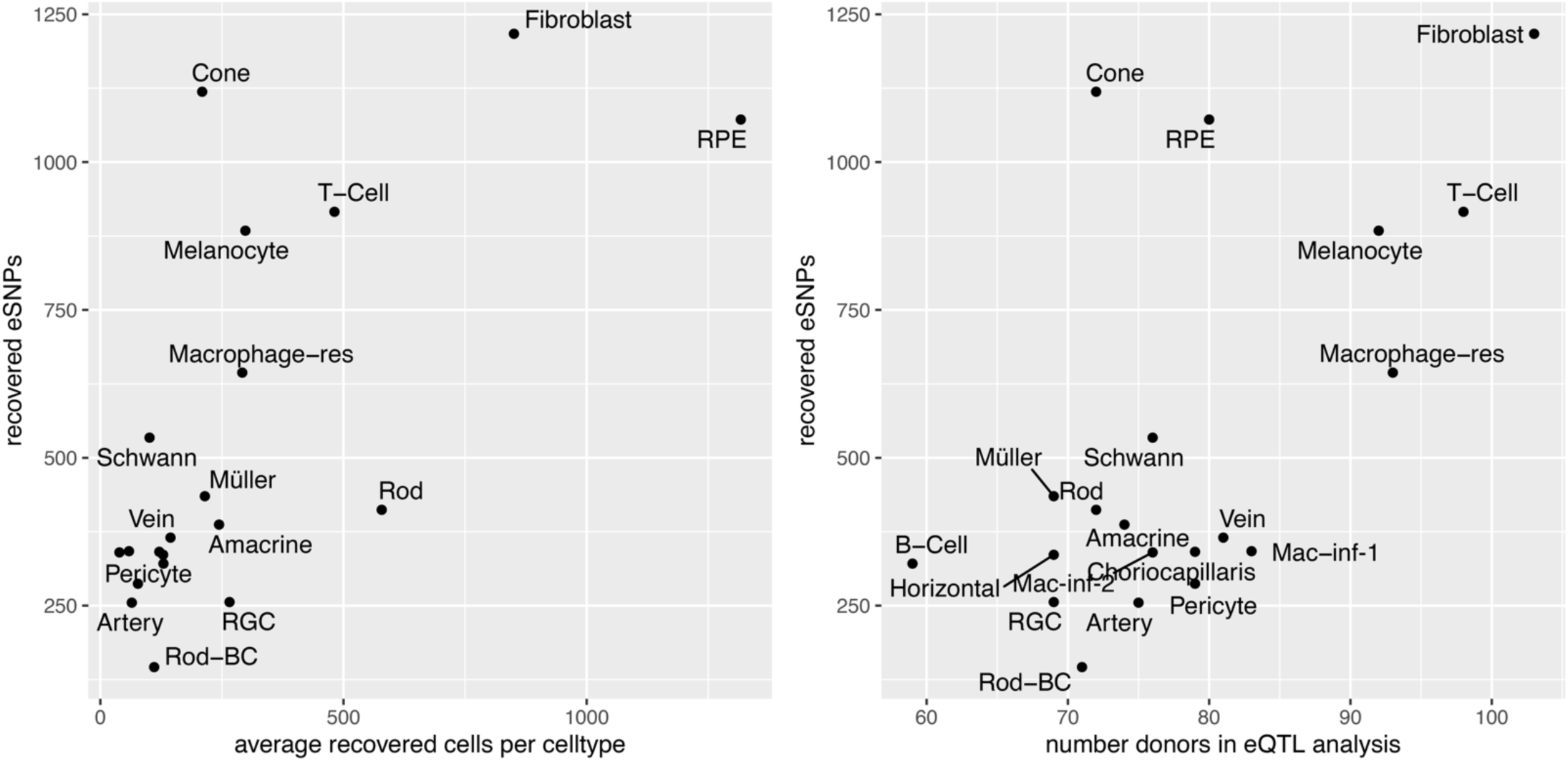
The number of significant eSNPs is related to the number of donors included in each eQTL analysis. **A.** The average number of cells recovered in each donor (x-axis) is compared to the number of eSNPs meeting genome wide significance (FDR < 0.05). More abundant cell types had more eSNPs meeting genome wide significance. **B.** The number of donors with at least 10 recovered cells in each cell type (x-axis) is compared to the number of eSNPs meeting genome wide significance (FDR < 0.05). Cell types with more donors in the eQTL analysis had more eSNPs meeting genome wide significance.

**SI Figure 3:**
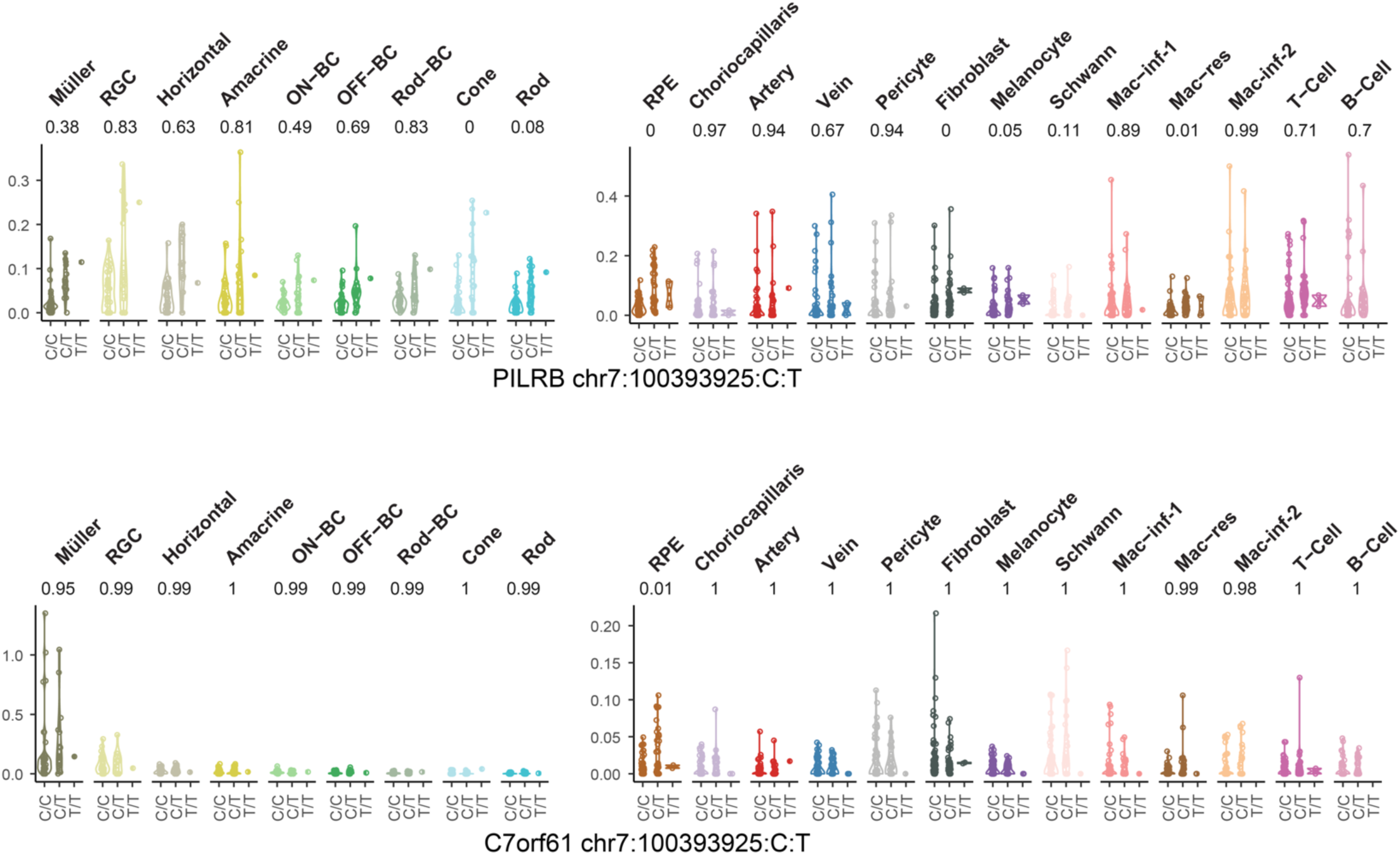
Expression of *PILRB* and *C7orf61* across rs6955367. **A.** Increasing number of risk alleles (T) at rs6955367 was associated with increased *PILRB* expression (**A**) as well as nearby *C7orf61* (*SPACDR*) expression (**B**). FDR < 0.05 significance was met in cone photoreceptors, RPE cells, fibroblasts for increased *PILRB* expression and RPE cells with increased *C7orf61* (*SPACDR*) expression.

**SI Figure 4:**
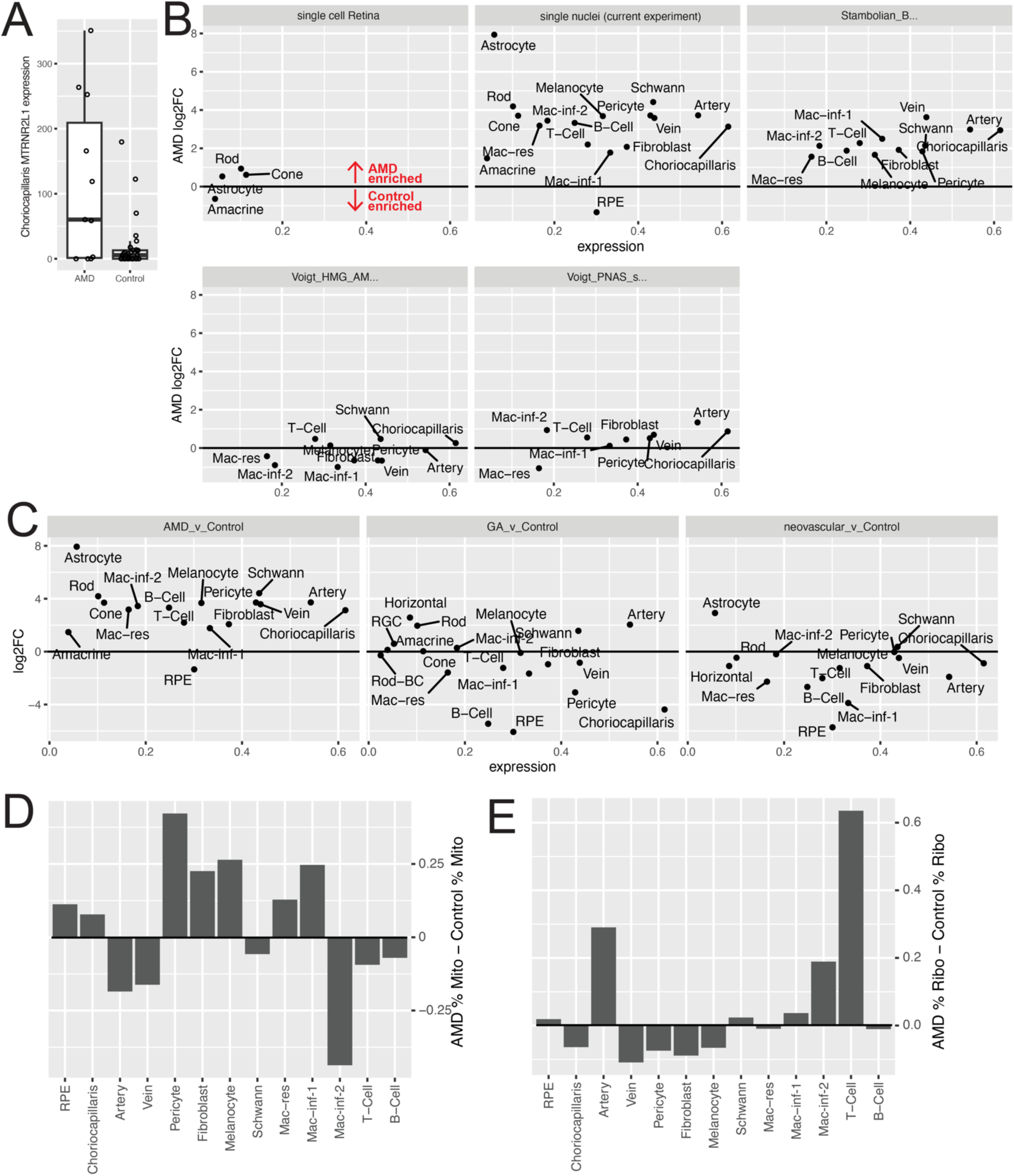
**A.** *MTRNR2L1* expression in the choriocapillaris split by AMD status. Each dot represents mean MTRNR2L1 expression for one donor and boxplots represent the mean and interquartile range. **B.** *MTRNR2L1* AMD enrichment across each cell type in the current snRNA-seq study as well as previously published studies with both AMD and control samples. The log2FC (y-axis) and cell type expression level (x-axis) of *MTRNR2L1* are displayed. *MTRNR2L1* is upregulated in AMD across multiple experiments. **C.** Comparison of differential expression results for dry AMD versus control, GA versus control, and MNV versus control. *MTRNR2L1* is not consistently enriched in MNV or GA comparisons. **D-E**. Differences in mitochondrial (D) and ribosomal (E) gene content between AMD and control samples. There is no clear trend of AMD samples having higher mitochondrial or ribosomal gene expression across all cell types, decreasing the likelihood that AMD samples have more contamination of ambient RNA.

